# CYSTM1, a drought-responsive cysteine-rich transmembrane module protein, negatively affects drought tolerance in *Arabidopsis thaliana*

**DOI:** 10.64898/2026.10.16.759560

**Authors:** Yasira Shoaib, Linus Geyer, Lena Berresheim, Julia Werdin, Igancio David Díaz Hernández, Carla Rack, Birte Brehm, Ute C. Vothknecht, Fatima Chigri

**Affiliations:** Institute for Cellular and Molecular Botany (IZMB), University of Bonn, Kirschallee 1, 53115 Bonn, Germany

**Keywords:** drought stress, *CYSTM1*, abscisic acid, transcriptomics, co-expression analysis

## Abstract

Drought stress represents a major challenge for plant growth and survival, requiring coordinated regulation of physiological and molecular responses to maintain water balance. The cysteine-rich transmembrane module (CYSTM) family comprises a highly conserved group of small proteins whose functions in plant stress adaptation remain largely unknown. Here, we identified *CYSTM1* (AT1G05340) as a drought-responsive gene in *Arabidopsis thaliana* whose expression increased in response to prolonged water deficit, osmotic stress, high salinity, and abscisic acid (ABA). Analysis of a loss-of-function T-DNA insertion mutant (*cystm1*) showed no growth phenotype compared to the wild-type under well-watered conditions. However, the mutant plants exhibited enhanced drought tolerance, whereas independent complementation lines restored the wild-type phenotype. The improved drought performance of the *cystm1* plants was associated with reduced stomatal aperture and decreased transpiration. Mutant plants also displayed an enhanced seedling establishment under osmotic stress conditions. Loss of *CYSTM1* altered ABA-associated responses at physiological and transcriptional level, resulting in reduced sensitivity to exogenous ABA and attenuated expression of ABA- and drought-responsive genes during water deficit. Co-expression analysis showed that genes with expression pattern similar to *CYSTM1* include many genes related to drought response as well as other members of the *CYSTM* family. Further expression analyses revealed selective transcriptional reprogramming within the *CYSTM* family following the loss of *CYSTM1*, suggesting crosstalk or functional interdependence within the CYSTM gene family. Collectively, these findings identify CYSTM1 as a novel negative effector of drought tolerance.

**Main conclusion:** Loss of *CYSTM1* is associated with altered ABA-related responses and transcriptional coordination within the Arabidopsis *CYSTM* gene family and enhances plant survival under progressive drought.

## Introduction

Plants, as sessile organisms, are subjected to a wide range of biotic and abiotic stresses that limit their growth, development, and yield (Gupta et al. 2020). Among these, drought is one of the major environmental constraints impairing respiration, photosynthesis, stomatal movements, and overall plant performance (Lv et al. 2023; Qiao et al. 2024). Drought arises from prolonged reductions in water availability, resulting in a sustained decrease in soil water potential that disrupts cellular homeostasis and ultimately plant growth (Salehi-Lisar and Bakhshayeshan-Agdam 2016). To cope with water deficit, plants activate complex physiological, cellular and molecular responses that enable them to maintain water status and minimize stress-induced damage (Bray 1997; Ingram and Bartels 1996; Haghpanah et al. 2024; Takahashi et al. 2020). These adaptive responses are accompanied by extensive transcriptional reprogramming, leading to the activation of genes whose gene products confer stress tolerance (Seki et al. 2002; Shinozaki and Yamaguchi-Shinozaki 2007; Maestro-Gaitán et al. 2025; Swift et al. 2026). A major component of this regulatory network is the phytohormone abscisic acid (ABA) that coordinates numerous physiological and transcriptional responses to drought (Iuchi et al. 2001; Muhammad Aslam et al. 2022). However, drought adaptation also involves ABA-independent signalling pathways highlighting the complexity of transcriptional networks that regulate the plant response to water deficit (Sakuma et al. 2006; Liu et al. 2018; Yoshida et al. 2014).

Under drought stress, ABA biosynthesis is rapidly induced, particularly in roots, followed by systemic transport to aerial organs where it coordinates transcriptional and physiological responses that promote stomatal closure and thus water conservation (Hu et al. 2016). ABA signalling operates through a highly conserved receptor–phosphatase–kinase module. ABA binds to members of the PYR/PYL/RCAR receptor family, resulting in inhibition of clade A PP2C phosphatases (Yoshida et al. 2010). This inhibition permits activation of SNF1-related kinase 2 (SnRK2s) proteins, which subsequently phosphorylate downstream transcription factors such as AREB/ABFs (Fujita et al. 2013). These transcriptional factors induce the spatial and temporal activation of numerous ABA-dependent drought responsive genes such as *RD29A* and *COR15A* that enhance osmoprotection, membrane stability, and cellular tolerance to water deficit (Narusaka et al. 2003; Meng et al. 2015).

Plant’s cellular and molecular responses to drought are ultimately translated into various physiological adjustments that enable plants to maintain water status and survive prolonged periods of water deficit. Among these adaptations, regulation of stomatal aperture, density and size are one of the most effective mechanisms for optimizing water-use efficiency (Basu et al. 2016). ABA-mediated signalling directly modulates guard cell ion channels, leading to efflux of potassium and anions, resulting in reduced guard cell turgor and thus stomatal closure (Kashtoh and Baek 2021). However, prolonged stomata closure restricts CO_2_ uptake, leading to reduced photosynthetic performance and growth. Consequently, precise regulation of stomatal behaviour is essential to balance water conservation with carbon acquisition, making ABA a critical integrator linking molecular signalling cascades to physiological adaptation to drought. Drought also induces extensive changes in root system architecture which can facilitate water acquisition under water-limited conditions. While primary root elongation is often sustained or even promoted to facilitate access to deeper soil moisture, lateral root initiation and outgrowth are often suppressed under water limitation (Deak and Malamy 2005). These developmental adjustments operate in concert with physiological responses, forming an integrated adaptive framework that enables plants to balance water acquisition with water conservation during prolonged water deficit.

Cysteine-rich Transmembrane Module (CYSTM) Proteins constitute a family of small, non-secreted proteins characterized by a conserved cysteine-rich transmembrane domain near the C-terminus. (Venancio and Aravind 2010). CYSTM proteins are widely conserved across eukaryotes, including fungi, animals, and plants. Functional studies in plants and yeast have implicated several members of this family in responses to environmental stress (Venancio and Aravind 2010; Xu et al. 2018). In *Arabidopsis thaliana*, the CYSTM family consists of 13 paralogs distributed on different chromosomes and their promoter regions contain various cis-acting elements responsive to biotic stress (fungi), abiotic stresses (salt, drought, cold, heat), and hormones such as ABA, jasmonic acid (JA), gibberellic acid (GA), salicylic acid (SA), and ethylene (Xu et al. 2018). The CYSTM family is also present in other plants species such as in *<u>D</u>igitaria ciliary* and *Oryza sativa (rice), Capsicum annuum L* (pepper)*, Beta vulgaris* (sugar beet) and *Canavalia rosea* (Kuramata et al. 2009; Ding et al. 2023; Ghaemi et al. 2020). Despite this evolutionary conservation, the biological functions of most plant CYSTM proteins remain poorly understood. Nevertheless, emerging evidence indicates that individual family members may participate in various developmental and stress-related processes. For example a recent study in Chinese cabbage identified nine CYSTM members and demonstrated a role for *CYSTM1* in regulating leaf size development (Zheng et al. 2026).

RNA-seq analysis showed *CYSTM1* to be upregulated under progressive drought (Mahmud et al. 2022). Although *CYSTM1* in *Arabidopsis thaliana* has previously been implicated in responses to heat, UV and pathogen (Joshi et al. 2020; Pereira Mendes et al. 2021), any role in drought stress remains unknown. To address this question, we examined the contribution of *CYSTM1* to drought tolerance and ABA-associated responses using a T-DNA insertion mutant and two independent complementation lines. Our data show *CYSTM1* to be an ABA-mediated negative effector of drought tolerance, whose absence changes the expression of various ABA- and drought-responsive genes. Loss of *CYSTM1* also altered the expression of other CYSTM family members under control and drought stress conditions, suggesting crosstalk and/or functional interdependence within the CYSTM gene family.

## Methods

### Plant material and growth conditions

*Arabidopsis thaliana* ecotype Columbia (Col-0) was used as the wild-type in this study. The confirmed T-DNA insertion *cystm1* mutant line (SALK_124375C) was ordered from the Nottingham Arabidopsis Stock Centre (NASC, UK). Plants were grown under long-day conditions (18 h light / 6 h dark) at 22 °C either on ½ MS media supplemented with 1% (w/v) sucrose and 6 % (w/v) phytoagar or in soil. Illumination was provided at an intensity of 100 μmol m⁻² s⁻¹ using Philips TLD 18W fluorescent lamps with alternating 830/840 light spectra.

### Mutant confirmation and generation of stable transgenic Arabidopsis lines

For the *cystm1* mutant line, the T-DNA insertion in the 2^nd^ exon was confirmed through genotyping PCR using two set of primer pairs (Supplementary Figure S1a and b) using forward and reverse primers flanking the T-DNA insertion mutation, as suggested by Signal Salk website (http://signal.salk.edu/cgi-bin/tdnaexpress). In addition, the reverse primer was combined with the T-DNA specific LBb1.3 primer. For the generation of complementation lines expressing 35S::*CYSTM1*-YFP construct, the coding sequence of *CYSTM1* (without stop codon) was cloned into the pBIN-19 ANX vector in frame with the coding sequence of the YFP reporter gene. The expression cassette transformed into the *cystm1* mutant genome using *Agrobacterium tumefaciens* and floral dipping (Zhang et al. 2006). Two independent T3 transgenic lines were selected through glufosinate resistance and constitutive expression was confirmed through RT-qPCR (Supplementary Figure S2c and d). The primer sequences used for cloning and screening are listed in Supplementary Table S1.

### Plant phenotyping in soil and measurement of relative water content (RWC %)

For phenotyping under normal and drought stress conditions, the seeds of WT, *cystm1* and two independent complementation lines were germinated in batches and after 7 days were transferred to individual pots containing 100 grams of potting soil. Plants were watered regularly until day 18 after germination, when all the pots were weighted and set to 200 grams by adding water. Watering was then stopped for progressive drought conditions and pictures were taken every day after 10 days of water withholding. Pots of control and drought exposed plants were randomized every day to avoid positional effect on plant growth. The relative leaf water content (RWC %) was measured following previously published protocol (Shoaib et al. 2026) by measuring three different weights, i.e Fresh weight (FW) measured immediately after removing the leaves from plants, turgid weight (TW) after submerging leaves in water for 24 hours, and dry weight (DW) after drying the leaves at 70 °C for 72 hours. The RWC (%) was calculated using the formula (FW-DW) / (TW-DW) × 100.

### Stomatal aperture, density and transpiration measurements

For the determination of stomatal aperture, 30-day-old plants of WT, *cystm1* mutant, and complementation lines grown under well-watered conditions (control) were incubated in imaging buffer (10 mM MES, 5 mM KCl, 50 μM CaCl_2_, pH 6.15) for two hours. The lower epidermis from the 6^th^ and 7^th^ leaves was carefully peeled, separated from mesophyll cells, and observed using the bright field setting of a Leica SP8 Lightning (Leica Biosystems, Germany). Pictures were taken using the internal LAS X software and stomatal aperture was measured using ImageJ (Schindelin et al. 2012). Stomatal density was determined as described (Bhattacharyya et al. 2025) using data from six individual plants for each genotype. Stomatal transpiration was measured with a LI-COR (LI-6000) porometer/fluorometer (LI-COR Environmental GmbH, Germany), using data from two leaves each of ten independent plants of each genotype.

### Phenotyping on MS media

Sterilized seeds were sown on ½ MS medium supplemented with mannitol or ABA as indicated and grown under long-day growth conditions. Seeds were considered germinated when the radical protruded out of the seed coat and germination rate was scored daily until all the seeds were germinated (Shoaib et al. 2021). To analyze the post-germination growth effect, cotyledon greening (%) and fresh weights were measured at day 9 and 14 after germination, respectively. For root growth analysis, the MS plates were placed in a vertical position and root length was measured using ImageJ software after 14 days of germination. All experiments were repeated at least three times.

### Gene regulation under abiotic stresses and ABA

To analyse the expression level of *CYSTM1* under abiotic stresses (PEG, mannitol and salt) and ABA, WT seedlings were grown on ½ MS media for 21 days and treated with PEG6000, mannitol, NaCl or ABA as indicated for 3, 6 and 24 hours. At the end of the treatment, 27 seedlings per biological replicate were frozen in liquid Nitrogen followed by RNA extraction and the expression level quantification of *CYSTM1* through RT-qPCR.

### *CYSTM1* co-expression network and GO enrichment analysis

A list of *CYSTM1* co-expressed genes was retrieved from ATTED-II ver.13.0 (https://atted.jp/) using the CoExSearch function (Obayashi et al. 2022). The co-expression network was generated using STRING ver.11.0 (https://version-11-0b.string-db.org/) by pasting the first 100 co-expressed genes from the ATTED-II list into the multiprotein search function and selecting *Arabidopsis thaliana* as the organism. The setting that hides disconnected nodes was chosen before network visualization. Gene Ontology (GO) enrichment analysis of the co-expressed gene set was performed using ShinyGO ver. 8.6 to identify significantly enriched biological processes (Ge et al. 2020). Genes with strong connection to drought response were selected to check their expression through RT-qPCR. The primers used for RT-qPCR are listed in Supplementary Table 1 and co-expressed genes with their functional annotations are listed in Supplementary Table 2.

### RNA extraction, cDNA synthesis and RT-qPCR

For RNA extraction, the plant material was grinded in liquid nitrogen using pre-cooled pestle and mortar. RNA was extracted from approximately 100 mg fine powder using Roboklon Plant RNA kit (Roboklon GmbH Berlin, Germany) according to the manufacturer’s instructions. The RNA quality was assessed by separating 3 µl of RNA on an 1% agarose gel, followed by its quantification using a Nanondrop one (Thermo Scientific). cDNA was synthesised from 1-1.5 µg of total RNA using the RevertAid first strand cDNA kit (Thermo Fisher Scientific, Deutschland) with OligodT-_18_ and random hexamer primers according to the manufacturer’s instructions. The reaction was carried out at 42 °C for 2 hours and terminated through a 5 min incubation at 75°C. RT-qPCR was carried out in a 96 well plate using the SYBR Green RT-PCR kit (Thermo Fisher Scientific, Germany) and a Bio-Rad CFX 96 system (Bio-Rad laboratories, Germany). The expression level was normalised to the geomean of *ACTIN2* and *TUBULIN2* and calculated using 2^^-ΔΔCT^ method (Vandesompele et al. 2002). All primers used for RT-qPCR are listed in Supplementary Table 1.

### Statistical Significance

Unless otherwise stated, datasets were evaluated using two-way ANOVA, followed by Tukey’s multiple comparisons test. Differences were considered statistically significant at *P* < 0.05. Number of biological replicates and, where applicable plants per replicate, are stated in the figures. The *CYSTM1* co-expression network was generated through STRING using the interaction score of 0.400 confidence. For Gene Ontology (GO) enrichment analysis, statistical significance was assessed using false discovery rate (FDR)-adjusted p-values calculated by ShinyGO, with an FDR cutoff of 0.05.

## Results

### Loss of *CYSTM1* prolongs survival under progressive drought

To gain insight into the role of *CYSTM1* during drought stress, we examined transcriptome data generated in our laboratory from Arabidopsis plants exposed to progressive water deficit (Mahmud et al. 2022). Within this dataset, *CYSTM1* showed one of the strongest increases in transcript abundance after 14 days of water-withholding, making it a promising candidate for further functional characterization. RT-qPCR analysis confirmed that *CYSTM1* transcript levels increased progressively during continuous drought (Fig. 1a). A first significant increase of expression can be observed as early as day 6 of water deprivation, increasing exponentially after 12 days, suggesting an association of *CYSTM1* with drought stress responses.

**Fig. 1.**
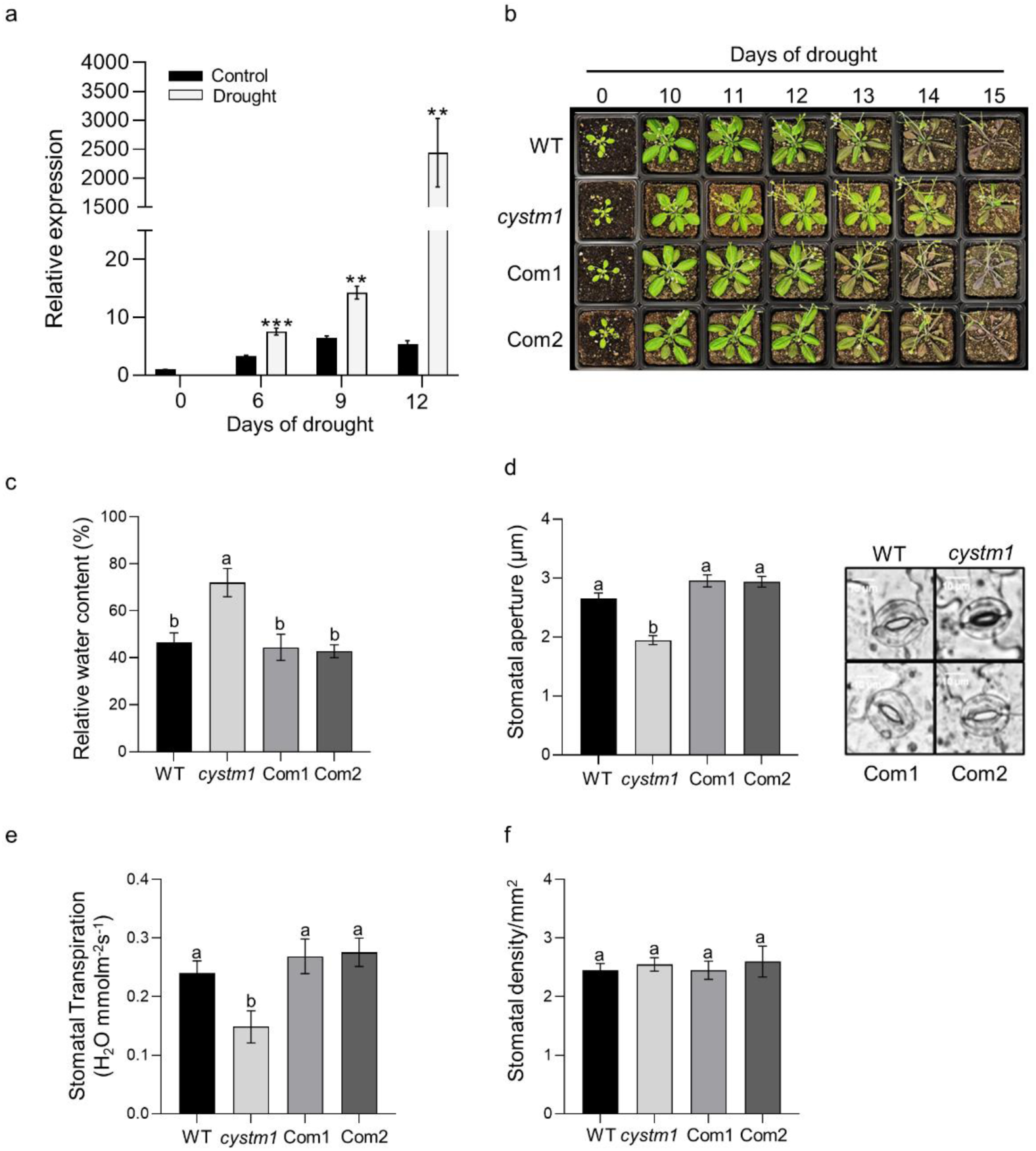
Loss of *CYSTM1* enhances drought tolerance by reducing stomatal-mediated water loss. (**a**) Relative transcript abundance of *CYSTM1* during progressive drought stress in 18-day-old wild-type (WT) plants grown in soil. Expression was normalized to the 0-day well-watered control. Values represent the mean ± SEM of three biological replicates. Data were analysed using Student’s t-test (P < 0.05). (**b**) Phenotypic response of WT, the *cystm1* mutant, and two independent complementation lines (Com1 and Com2) following progressive drought stress applied to 18-day-old soil-grown plants. (**c)** Relative water content (%) after 12 days of drought stress. Values represent the mean ± SE of six biological replicates (n = 6). Stomatal (**d**) aperture (n = 60), (**e**) transpiration (n = 20), and (**f**) stomatal density (n = 20) were measured in 30-day-old well-watered plants. Data were analysed using one-way ANOVA followed by Tukey’s multiple comparison test (P < 0.05)

To investigate the physiological role of CYSTM1 during drought stress, we characterized a confirmed Arabidopsis T-DNA insertion mutant (*cystm1*; SALK_124375C) together with two independent complementation lines expressing 35S::*CYSTM1*-YFP in the mutant background (Supplementary Fig. S1, a-c). RT-qPCR analysis confirmed the absence of *CYSTM1* transcripts in the mutant and restored expression in both complementation lines (Supplementary Fig. S1d). Under well-watered conditions, neither the mutant nor the complementation lines displayed obvious differences in vegetative or reproductive development compared to the wild-type (Supplementary Fig. S2a-g), indicating that CYSTM1 does not play a major role in normal plant growth and development.

To determine whether CYSTM1 contributes to drought adaptation, plants from the different lines were subjected to progressive water deprivation (Fig. 1b). Whereas wild-type and complementation plants exhibited first signs of wilting on day 12 and severe wilting after 14 days of drought, the *cystm1* mutant remained visibly less affected. This improved performance was accompanied by significantly higher relative water content (RWC) compared with wild-type and complemented plants (Fig. 1c), suggesting that lack of *CYSTM1* improves water retention during prolonged water deficit.

Since stomatal regulation is a major determinant of plant water status under drought, we next examined stomatal characteristics in the different genotypes. The *cystm1* mutant exhibited significantly narrower stomatal apertures than wild-type and the complementation lines (Fig. 1d), which was associated with a corresponding reduction in transpiration rate (Fig. 1e). In contrast, stomatal density did not differ between genotypes (Fig. 1f), indicating that altered stomatal regulation rather than changes in stomatal development contributes to the improved drought tolerance of the mutant.

### Transcriptional regulation of *CYSTM1* in response to osmotic, salt, and ABA treatments

We next determined whether *CYSTM1* responds to treatment with mannitol, polyethylene glycol (PEG), and NaCl, all of which produce osmotic or osmotic/ionic stress. We observed an increase in *CYSTM1* expression with all three treatments (Figure 2a-c), albeit with different kinetics. Whereas the response to PEG and NaCl peaked after 6 h and clearly declined by 24 h, the induction by mannitol increased steadily over the 24-h observation period. Moreover, in case of mannitol and NaCl, only the higher concentrations, i.e. 400 vs 300 mM mannitol and 200 vs 150 mM NaCl, resulted in pronounced induction, indicating that severe osmotic stress is required to elicit a substantial *CYSTM1* response.

**Fig. 2.**
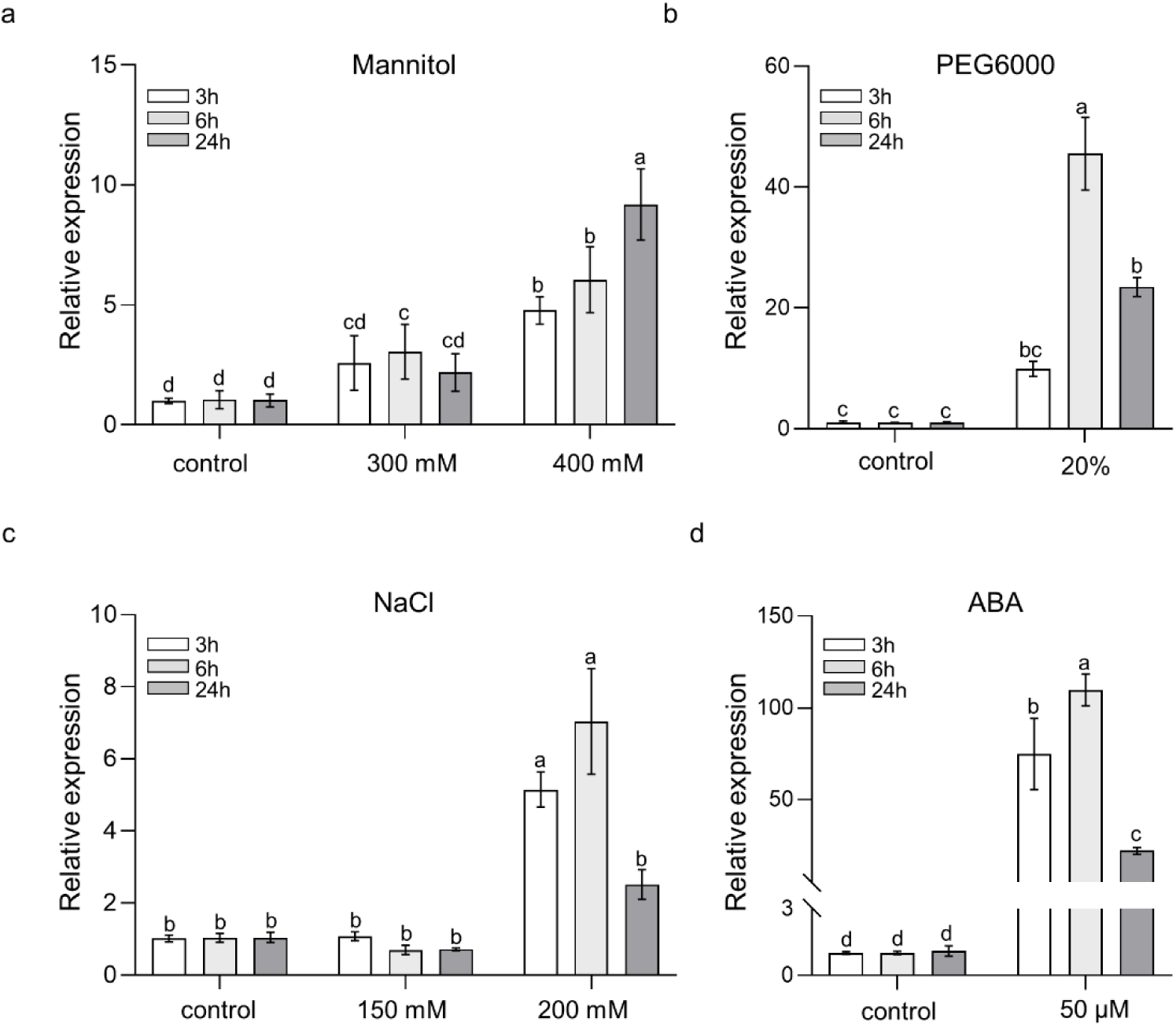
*CYSTM1* is responsive to various abiotic stresses and ABA. Relative transcript abundance of *CYSTM1* determined by RT-qPCR in 21-day-old wild-type (WT) seedlings grown on half-strength (½) MS medium following treatment with (**a**) 300 and 400 mM mannitol, (**b**) 20% PEG6000, (**c**) 150 and 200 mM NaCl, and (**d**) 50 µM ABA for 3, 6, and 24 h. Values represent the mean ± SE of six biological replicates (n = 6). Data were analysed using two-way ANOVA followed by Tukey’s multiple comparison test (P < 0.05)

Because abscisic acid (ABA) is a central regulator of plant drought responses, we further examined whether *CYSTM1* is responsive to ABA. Exogenous ABA treatment strongly induced its expression, with transcript levels reaching a maximum after 6 h (Fig. 2d), supporting a potential association of *CYSTM1* with ABA-mediated drought responses.

### *CYSTM1* contributes negatively to seedling establishment and root growth under non-favourable conditions

Given the induction of *CYSTM1* by drought and ABA, we next investigated whether *CYSTM1* contributes to stress adaptation during early plant development. To that end wild-type and *cystm1* mutant seedlings were grown on ½ MS medium supplemented with mannitol or ABA and their germination and post-germination growth responses were evaluated. (Fig. 3). Under control conditions, wild-type and *cystm1* seedlings exhibited comparable germination rates and early seedling development. Osmotic stress induced by 300 mM mannitol delayed germination by 1-2 days (Fig. 3a). The effect is similar to a treatment with ABA. However, no statistically significant differences were observed between wild-type and *cystm1* mutant, even though the mutant plants appeared to germinate a bit better under mannitol. The difference between wild-type and mutant became clearer in the post-germination analysis under mannitol (Fig. 3b). At day 9 after germination, the *cystm1* mutant exhibited a significantly higher proportion of seedlings with green cotyledons compared to wild-type plants and after 14 days showed a significantly greater fresh weight.

**Fig. 3.**
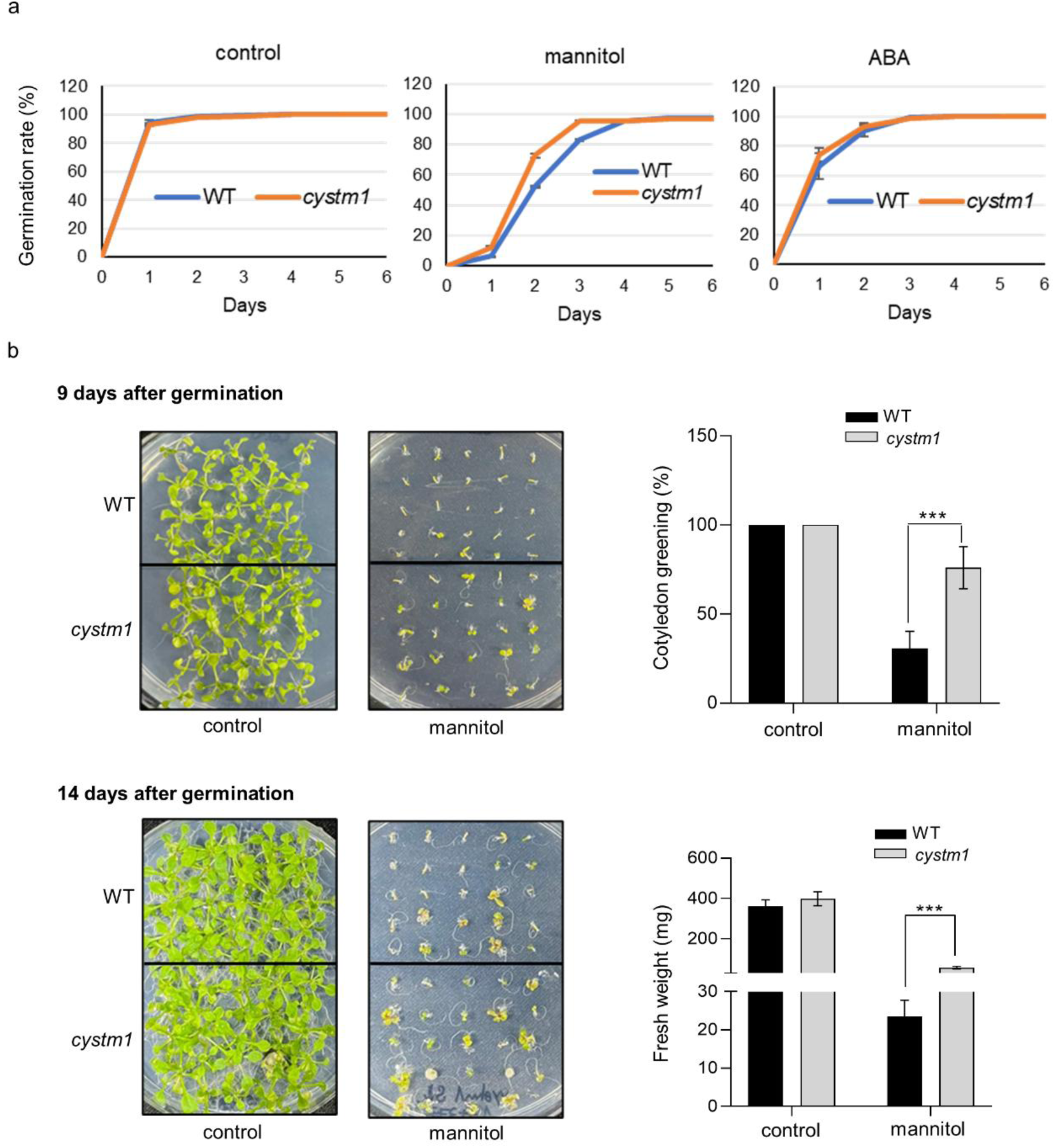
Loss of *CYSTM1* affects seedling establishment under non-favourable conditions. (**a**) Germination rate (%) of wild-type (WT) and *cystm1* seeds grown on half-strength (½) MS medium supplemented with H₂O (control), 0.5 µM ABA, or 300 mM mannitol (**b**) Representative images (left) and quantification (right) of cotyledon greening and (**c**) fresh weight of WT and *cystm1* seedlings grown on ½ MS medium supplemented with H₂O (control) or 300 mM mannitol for 9 days and 14 days, respectively. Values represent the mean ± SE of four biological replicates each 25 seedlings (n = 100). Data were analysed using multiple Student’s t-tests (P < 0.05)

Loss of *CYSTM1* also reduced the negative effect of mannitol on root elongation (Fig. 4a). Similarly, root elongation of the *cystm1* mutant was significantly less affected by ABA treatment compared to wild-type plants at both 0.5 and 1 µM ABA concentrations (Fig. 4b). This indicates that loss of *CYSTM1* reduces ABA sensitivity during root development, which likely contributes to the enhanced root growth of the mutant in the presence of mannitol.

**Fig. 4.**
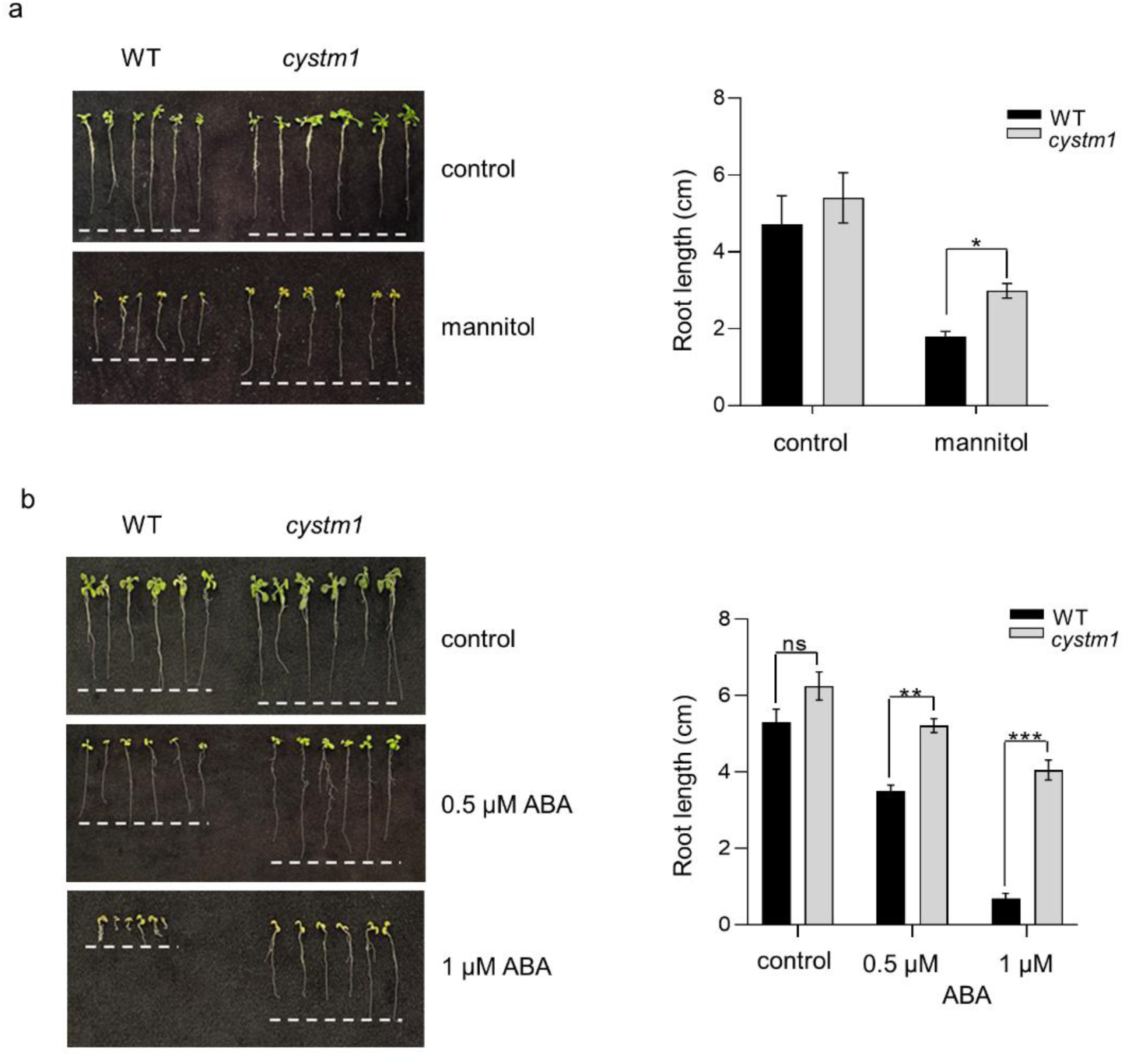
Loss of *CYSTM1* affects root growth under mannitol and ABA. Representative images (left) and root length measurements (right) of WT and *cystm1* seedlings grown on ½ MS medium supplemented with (**a**) 300 mM mannitol and (**b**) 0.5 and 1 µM ABA for 14 days. The dashed line serves as a visual reference for comparing root lengths among genotypes and treatments. Values represent the mean ± SE of four biological replicates each five seedlings (n = 20). Data were analysed using multiple Student’s t-tests (P < 0.05)

### Loss of *CYSTM1* affects the expression of various drought-responsive genes

To further investigate whether the altered physiological responses of *cystm1* were accompanied by changes in ABA-related gene expression, transcript levels of representative genes involved in ABA biosynthesis, signalling, and ABA-dependent drought responses were analysed in soil grown plants under progressive drought (Fig. 5).

**Fig. 5.**
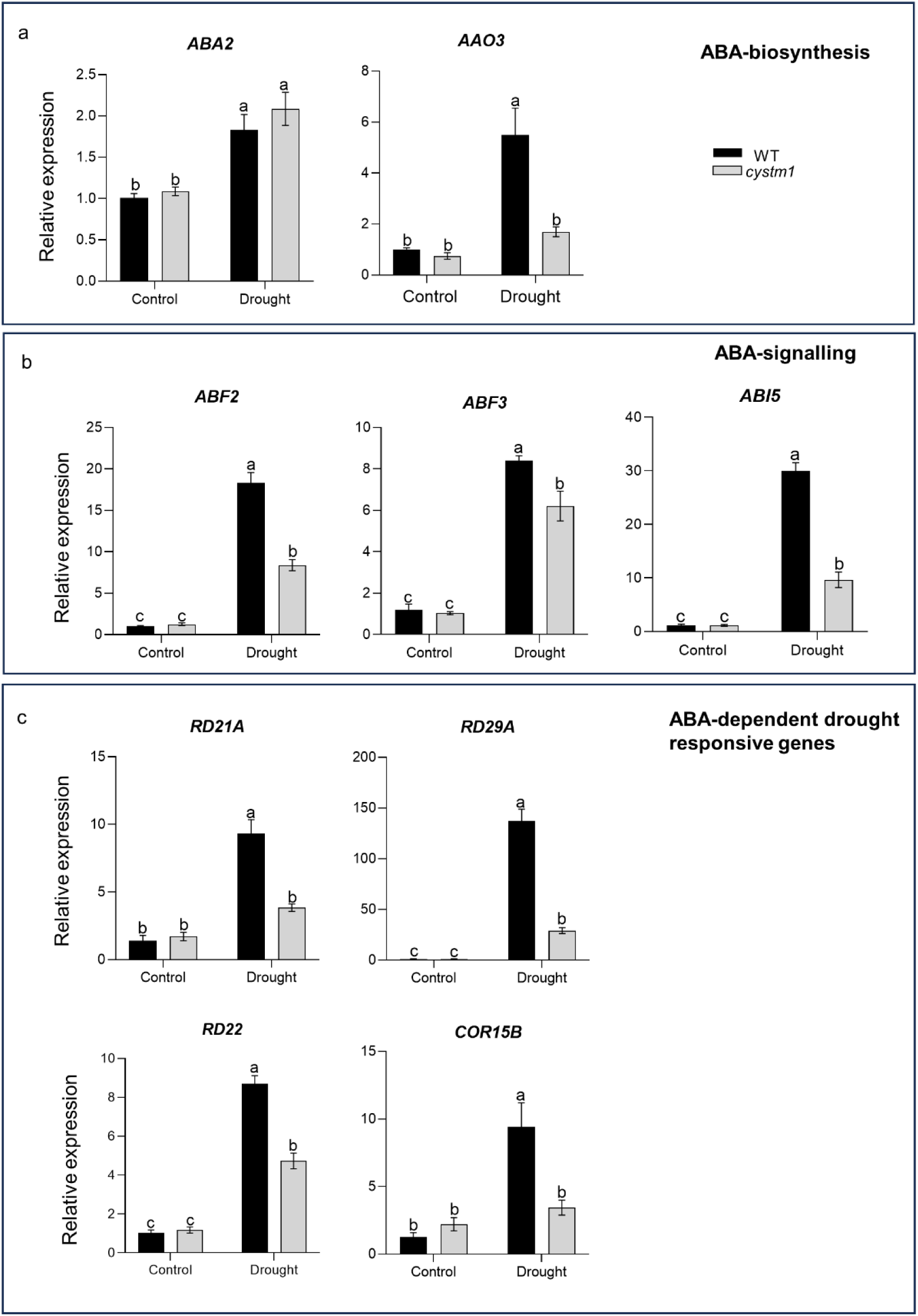
Relative transcript abundance of ABA-related genes. Relative transcript abundance of (**a**) ABA biosynthesis-, (**b**) ABA signalling-, and (**c**) ABA-dependent drought-responsive genes was determined by RT-qPCR in wild-type (WT) and *cystm1* plants under well-watered conditions and after 12 days of drought stress. Values represent the mean ± SE of three biological replicates (n = 3). Data were analysed using two-way ANOVA followed by Tukey’s multiple comparison test (P < 0.05)

While expression of the ABA biosynthetic gene *ABA DEFICIENT 2* (*ABA2*) was comparable between the two genotypes under control and drought conditions, drought-induction of *ABSCISIC ALDEHYDE OXIDASE 3* (*AAO3*) was significantly increased in wild-type plants but remained largely unchanged in the *cystm1* mutant (Fig. 5a). Similarly, several transcriptional regulators involved in ABA signalling, such as *ABF2*, *ABF3*, and *ABI5*, showed reduced transcript accumulation in the *cystm1* mutant compared to wild-type plants under drought stress (Fig. 5b), as well as ABA- and drought-responsive genes such as *RD21A*, *RD29A*, *RD22*, and *COR15B* (Fig. 5c). Together, these findings suggest that loss of *CYSTM1* attenuates ABA responses at both physiological and transcriptional levels.

Genes involved in related biological processes frequently display similar expression patterns. We therefore identified genes co-expressed with *CYSTM1* using the ATTED-II database and selected the top 100 genes based on their co-expression scores (Supplementary Table S2). These genes were used to construct an interaction network which connects *CYSTM1* to a cluster enriched in drought- and dehydration-responsive genes (Figure 6a). Furthermore, GO analysis of these genes showed enrichment of biological processes such as responses to water deprivation, ABA, and several other biotic and abiotic stresses (Figure 6b), supporting an association of *CYSTM1* with stress-responsive pathways.

**Fig. 6.**
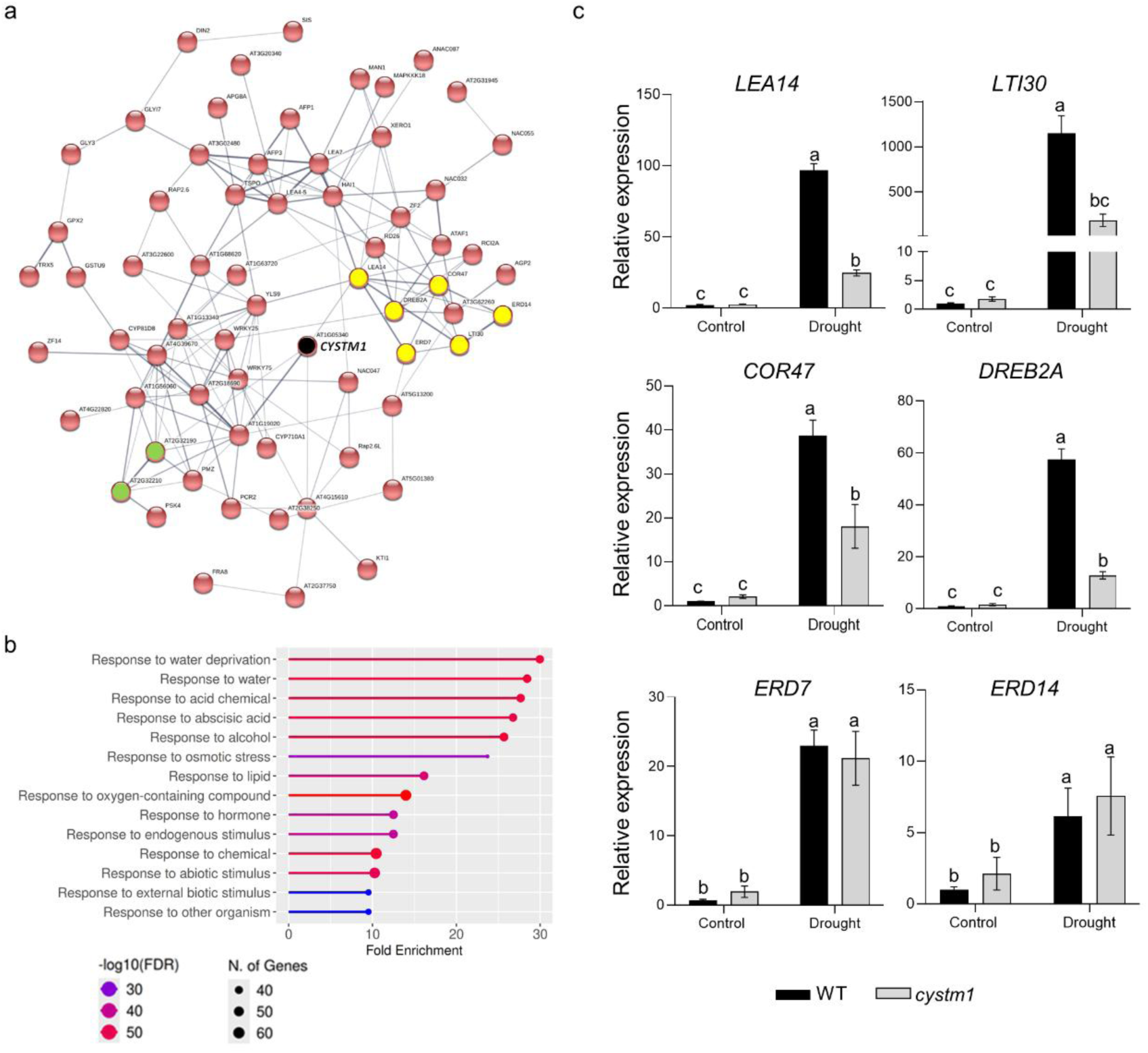
CYSTM1-associated gene network and functional enrichment analysis. (**a**) STRING interaction network of the 100 genes most strongly co-expressed with CYSTM1 (AT1G05340, black-filled circle), identified based on ATTED-II co-expression scores. Green-filled circles: *CYSTM4* (AT2G32190) and *CYSTM6* (AT2G32210); yellow-filled circles: drought-responsive genes selected for validation using RT-qPCR. (**b**) Gene Ontology (GO) enrichment analysis of the 100 *CYSTM1*-associated genes performed using ShinyGO. (**c**) Relative transcript abundance of selected drought-responsive genes determined by RT-qPCR. Values represent the mean ± SE of three biological replicates. Data were analysed using two-way ANOVA followed by Tukey’s multiple comparison test (P < 0.05)

Based on their established roles in drought responses, a subset of genes from the co-expression network, i.e. *LEA14*, *COR47*, *LTI30*, *DREB2A*, *ERD7*, and *ERD14,* were selected for subsequent validation using RT-qPCR. Under progressive drought, the expression of all selected genes increased in both genotypes, however, the transcript levels of *LEA14*, *COR47*, *LTI30*, and *DREB2A*, were significantly higher in wild-type plants than in the *cystm1* mutant (Fig. 6c). By contrast, the expression of two early drought-responsive genes, *ERD7* and *ERD14,* was largely unaffected in the *cystm1* mutant. These results indicate that *CYSTM1* modulates the expression of a subset of drought-responsive genes rather than broadly controlling drought-induced transcription.

### Loss of *CYSTM1* reshapes transcriptional regulation within the Arabidopsis CYSTM gene family

*CYSTM1* is part of small gene family and two of its members, *CYSTM4* (AT2G32190) and *CYSTM6* (AT2G32210), where found as part of the *CYSTM1*-associated co-expression network (Fig. 6a), suggesting potential regulatory relationships within the CYSTM gene family. To investigate whether loss of *CYSTM1* influences the expression of other members, transcript levels of the remaining 12 *CYSTM* genes were analysed in wild-type and *cystm1* plants.

Under well-watered conditions, transcript levels of *CYSTM2*, *CYSTM4*, *CYSTM5*, *CYSTM10*, *CYSTM11*, and *CYSTM12* were significantly higher in the *cystm1* mutant than in wild-type plants (Fig. 7), indicating that loss of *CYSTM1* affects the normal transcriptional balance of selected family members even in the absence of drought stress. Exposure to drought further accentuated these genotype-dependent differences. Of the 6 *CYSTM* genes already affected under well-watered conditions, all except *CYSTM5* also showed increased expression in the mutant under drought. In case of *CYSTM4* and *CYSTM12* this effect is much more pronounced than under control conditions. In addition, *CYSTM7*, *CYSTM9*, and *CYSTM13* showed up-regulation in the mutant only under drought conditions. The other *CYSTM* genes showed no or only minor differences under either well-watered or drought conditions. These include *CYSTM6*, despite the close co-expression relationship to *CYSTM1* predicted by the co-expression network. Collectively, these findings demonstrate that loss of *CYSTM1* selectively reshapes basal and drought-responsive expression within the Arabidopsis CYSTM gene family.

**Fig. 7.**
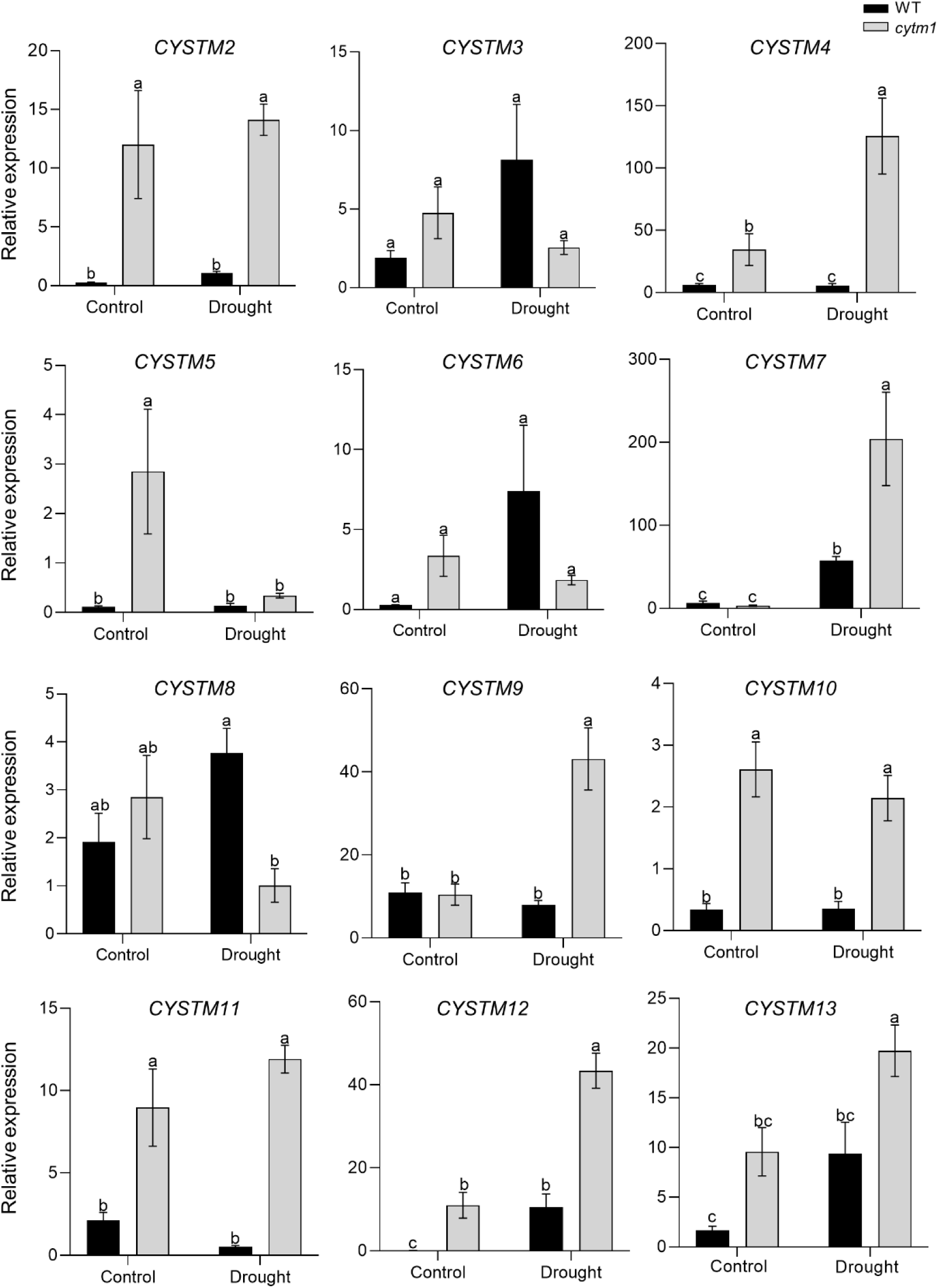
Loss of *CYSTM1* alters the expression of Arabidopsis *CYSTM* family genes. Relative transcript abundance was determined by RT-qPCR in wild-type (WT) and *cystm1* plants under well-watered conditions and after 12 days of drought. Values represent the mean ± SE of six biological replicates (n = 6). Data were analysed using two-way ANOVA followed by Tukey’s multiple comparison test (P < 0.05)

## Discussion

In the present study, we identified *CYSTM1* as a drought-responsive gene whose transcript abundance increased markedly during progressive water deficit. However, loss of *CYSTM1* enhanced drought tolerance, indicating that *CYSTM1* functions as a negative modulator of plant drought adaptation. While the strong induction of *CYSTM1* by drought may initially appear counterintuitive, loss of modulating components can enhance drought tolerance by releasing adaptive constraints on stress-responsive pathways (Wilson et al. 2009). These complex feedback mechanisms fine-tune the intensity and duration of stress responses, thereby balancing survival with growth (Yoshida et al. 2014).

The phytohormone ABA is a central regulator of plant responses to drought stress, coordinating physiological and transcriptional adaptations that enhance water conservation and stress tolerance (Haghpanah et al. 2024). The strong induction of *CYSTM1* by ABA poses the question whether CYSTM1 functions within the ABA-regulatory network. Indeed, loss of *CYSTM1* significantly attenuated physiological and transcriptional responses that are closely associated with this hormone. The *cystm1* mutant exhibited reduced sensitivity to exogenous ABA during primary root growth and showed diminished drought-induced expression of the core ABA signalling transcription factors *ABF2*, *ABF3*, and *ABI5*. Likewise, transcript levels of several ABA-responsive marker genes, including *RD21A*, *RD22*, *RD29A*, and *COR15B*, were also markedly reduced in the mutant under drought stress. Loss of *CYSTM1* also attenuated the induction of *LEA14*, *COR47*, *LTI30, and DREB2A*, genes that were found among the most strongly co-expressed genes with *CYSTM1*. They are widely recognized as important components of plant responses to dehydration and are typically induced to protect cellular structures and maintain osmotic homeostasis during water deficit (Yamaguchi-Shinozaki and Shinozaki 2006; Sakuma et al. 2006; Shi et al. 2015). GO enrichment analysis of the top 100 genes co-expressed with *CYSTM1* revealed significant enrichment for responses to water deprivation, ABA, and other biotic and abiotic stresses. This indicates that *CYSTM1* operates within a stress-responsive regulatory network integrating ABA-dependent and other drought-responsive pathways, a characteristic feature of multilayered stress signalling in plants (Yoshida et al. 2014; Margay et al. 2024). This interpretation is supported by studies of the Arabidopsis CYSTM family member *CYSTM3*, which acts as a negative regulator of abiotic stress tolerance through modulation of ion homeostasis and reactive oxygen species balance (Xu et al. 2019). Together, these observations suggest that negative regulation may represent a conserved functional characteristic of at least a subset of CYSTM family members in plants. On the other hand, we observed unchanged expression of *ERD7* and *ERD14*. Both genes have been described as early responders to dehydration (Kiyosue et al., 1994a, b), with their expression remaining elevated during prolonged dehydration. Thus, their expression levels at day 12 of drought may reflect an early-established stress response that is less sensitive to differences associated with the loss of *CYSTM1*. This interpretation is consistent with the relatively weak induction of *CYSTM1* at day 6 of drought. Moreover, the requirement for higher concentrations of mannitol and NaCl to induce *CYSTM1* expression on plates suggests that *CYSTM1* responds preferentially to more pronounced stress conditions.

Under progressive soil drying, *cystm1* plants maintained higher relative water content and exhibited reduced wilting compared with wild-type plants. The observation that stomatal aperture and thus transpiration rates were significantly reduced without changes in stomatal density, indicates that the improved drought tolerance of the mutant resulted from altered stomatal behaviour rather than developmental changes in stomatal patterning. Stomatal regulation represents a key adaptive mechanism that minimizes water loss, preserves cellular turgor, and contributes to drought tolerance under water-deficit conditions (Chen et al. 2024; Razzak et al. 2026). Importantly, improved physiological adaptation can lessen the severity of cellular dehydration, thereby reducing the requirement for extensive transcriptional reprogramming. In the *cystm1* mutant, enhanced water conservation through reduced transpiration and improved stomatal regulation is therefore likely to mitigate the physiological impact of drought, decreasing the demand for strong activation of downstream drought-responsive genes. This interpretation is consistent with studies demonstrating that the magnitude of drought-induced transcriptional responses closely reflects the severity of the physiological stress experienced by the plant (Swift et al. 2026).

Enhanced drought performance of the *cystm1* mutant was also evident during early seedling development. The absence of differences during seed germination, together with the pronounced effects observed during post-germination development, suggests that the role of *CYSTM1* in stress responses depends on the developmental context. Together with the enhanced survival of *cystm1* plants under progressive drought, these findings indicate that loss of *CYSTM1* affects stress responses at different stages of vegetative development. Importantly, primary root growth and cotyledon greening are not strongly associated with stomatal regulation, suggesting that *CYSTM1* either affects multiple drought-response processes or that the observed changes in stomatal regulation are an indirect consequence of its function.

Loss of *CYSTM1* also caused a broader reorganization of transcriptional regulation within the Arabidopsis CYSTM family. Rather than eliciting a uniform transcriptional response, disruption of *CYSTM1* differentially affected the basal expression and drought responsiveness of individual family members. The selective nature of these transcriptional changes further supports previous evidence that CYSTM family members have undergone functional specialization, with diversification of their regulatory regions potentially allowing differential responses to environmental and hormonal signals despite their highly conserved cysteine-rich transmembrane domain (Xu et al. 2018; Pereira Mendes et al. 2021; Cai et al. 2025). Such selective transcriptional reorganization is consistent with the concepts of subfunctionalization and genetic robustness, whereby duplicated genes retain partially overlapping functions while acquiring specialized regulatory properties that can increase the flexibility and resilience of stress responses (Gu et al. 2003; Moore and Purugganan 2005). In our experimental set-up, the only other family member showing drought-induced up-regulation in the wild-type was *CYSTM7*. This appears to differ from the findings of Xu et al. (2018). However, their study used PEG 6000 treatment of plate-grown plants, and PEG treatment and soil drought have been shown to elicit substantially different transcriptomic responses (Gonzalez et al. 2024). More importantly, several CYSTM genes, i.e. *CYSTM4, CYSTM9, CYSTM12,* and *CYSTM13* are not drought-induced in the wild-type but showed varying degrees of increased expression in the *cystm1* mutant. Remarkably, these family members neither belong to the same phylogenetic subgroup nor share the same subcellular localization (Xu et al., 2018). Moreover, *CYSTM4* and *CYSTM12* already showed moderately increased expression in the *cystm1* mutant under control conditions. These patterns may indicate compensatory transcriptional changes within the CYSTM family following loss of CYSTM1, which become more pronounced under drought stress. Whether these changes reflect direct functional or regulatory interactions or indirect consequences of altered stress responses remains to be determined.

Taken together, our findings expand the understanding of the biological functions of the largely unexplored CYSTM protein family, especially CYSTM1. Moreover, this study further highlights the importance of negative modulating mechanisms in shaping plant responses to adverse environments and provides a framework for future studies aimed at dissecting their molecular functions and exploring their potential for improving plant resilience to environmental stress.

## Supplementary Information

Supplementary file 1 (XLSX, Supplementary Table S1)

Supplementary file 2 (XLSX, Supplementary Table S2)

Supplementary file 3 (PDF, Supplementary Fig. S1 and Supplementary Fig. S2)

## Authors contribution statement

YS contributed to conceptualization, investigation (responsible for most experimental work), formal analysis (responsible for statistical analysis), validation, visualization, and writing - original draft as well as review & editing. LG, CR, IDH, LB, JW, BB contributed to investigation (gene expression, phenotyping, screening, and molecular cloning), UCV contributed to conceptualization, validation, funding acquisition, project administration, supervision, and writing - review & editing). FC contributed to conceptualization, validation, visualization, supervision, and writing - original draft as well as review & editing. All authors contributed to the article and approved the submitted version.

