## Supplementary material for "CYSTM1, a drought-responsive cysteine-rich transmembrane module protein, negatively affects drought tolerance in *Arabidopsis thaliana*": Table S1, Table S2 , Figure S1 and Figure S2: Supplementary File 3.pdf

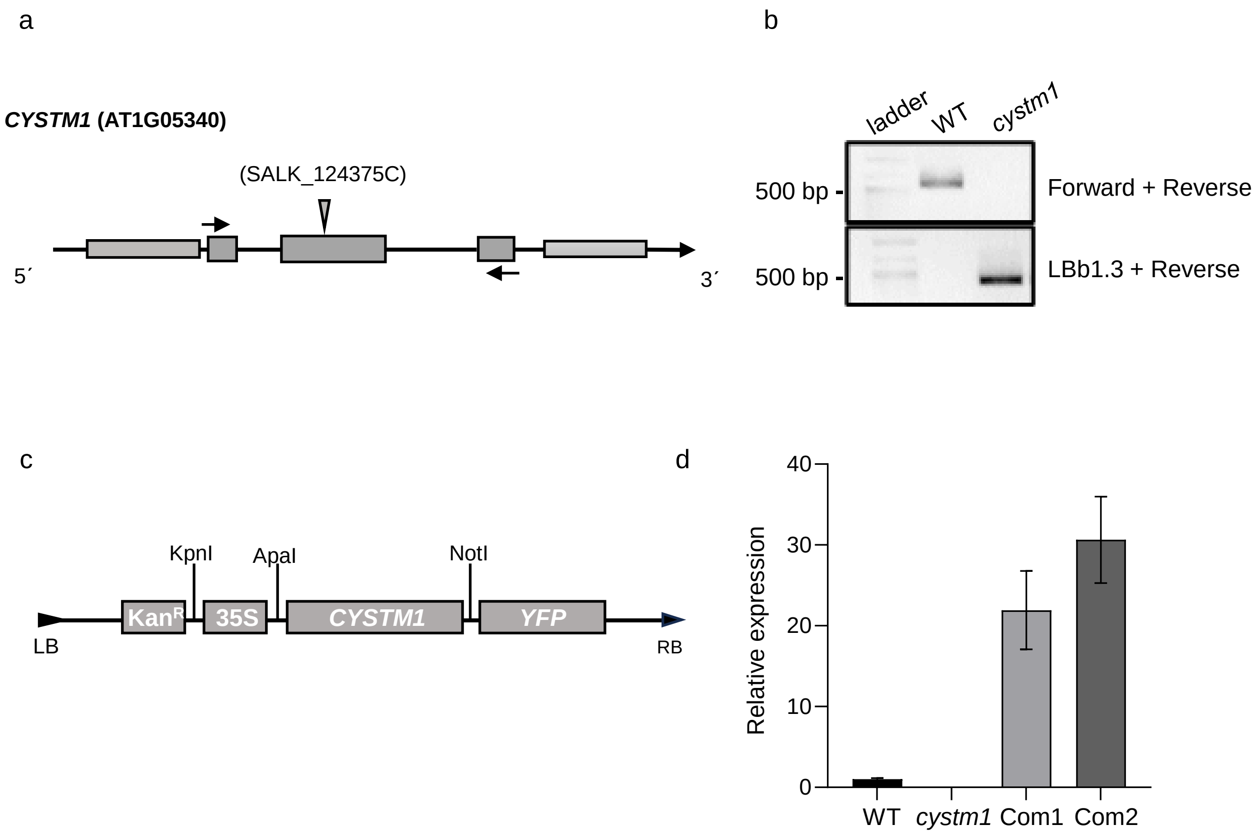


**Supplementary Fig. S1** Confirmation of the *cystm1* mutant and complementation lines. (**a**) Schematic representation of the T-DNA insertion site in the SALK_124375C line. Arrows indicate the positions of the forward and reverse primers used for genotyping PCR. Exons are represented by grey boxes and introns by grey lines. (**b**) Confirmation of a homozygous *cystm1* T-DNA insertion mutant by genotyping PCR. (**c**) Schematic representation of the complementation construct used to generate transgenic lines in the *cystm1* mutant background. (**d**) Confirmation of *CYSTM1* transcript abundance in the *cystm1* mutant and two independent complementation lines (Com1 and Com2) by RT-qPCR in rosette leaves of plants grown on soil under control conditions

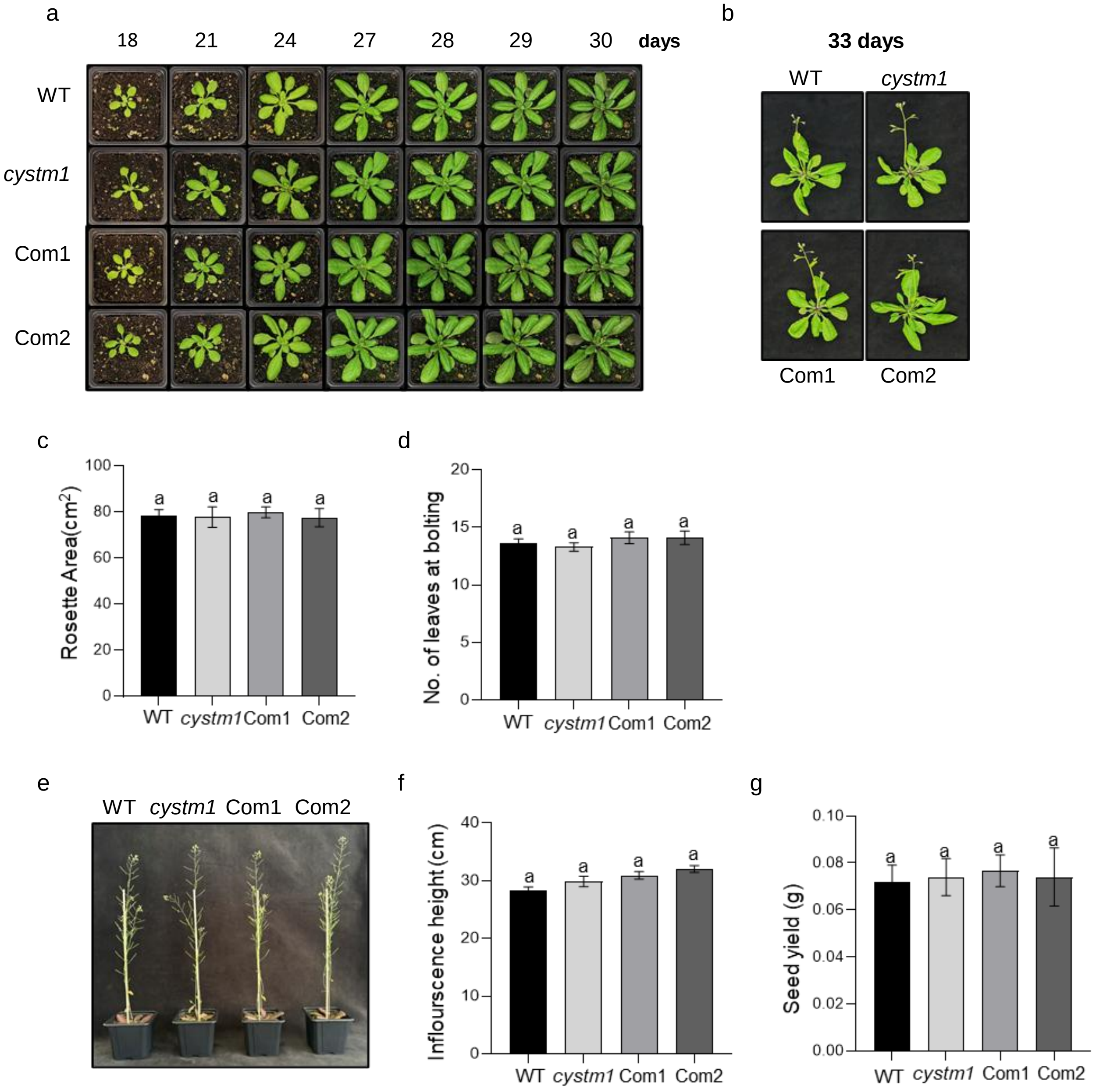


**Supplementary Fig. S2 Loss of *CYSTM1* does not affect plant growth and development under normal conditions.** (**a**) Representative images of whole rosettes at different time points after germination. (**b**) Flowering phenotype at 33 days after germination. (**c**) Rosette area measured at 30 days after germination. (**d**) Number of leaves at 30 days after germination. (**e**) Representative images of inflorescence development and (**f**) quantification of inflorescence height at 40 days after germination. (**g**) Seed yield under normal growth conditions. For all experiments the values represent means ± SE from 10 individual plants (n = 10). Statistical analysis was performed using one-way ANOVA followed by Tukey's multiple comparison test ($P < 0.05$)
